# Combining multiple functional connectivity methods to improve causal inferences

**DOI:** 10.1101/841890

**Authors:** Ruben Sanchez-Romero, Michael W. Cole

## Abstract

Cognition and behavior emerge from brain network interactions, suggesting that causal interactions should be central to the study of brain function. Yet approaches that characterize relationships among neural time series—functional connectivity (FC) methods—are dominated by methods that assess bivariate statistical associations rather than causal interactions. Such bivariate approaches result in substantial false positives since they do not account for confounders (common causes) among neural populations. A major reason for the dominance of methods such as bivariate Pearson correlation (with functional MRI) and coherence (with electrophysiological methods) may be their simplicity. Thus, we sought to identify an FC method that was both simple and improved causal inferences relative to the most popular methods. We started with partial correlation, showing with neural network simulations that this substantially improves causal inferences relative to bivariate correlation. However, the presence of colliders (common effects) in a network resulted in false positives with partial correlation, though this was not a problem for bivariate correlations. This led us to propose a new combined functional connectivity method (combinedFC) that incorporates simple bivariate and partial correlation FC measures to make more valid causal inferences than either alone. We release a toolbox for implementing this new combinedFC method to facilitate improvement of FC-based causal inferences. CombinedFC is a general method for functional connectivity and can be applied equally to resting-state and task-based paradigms.

## Introduction

A goal of brain connectivity research is to estimate mechanistic network architectures that define functional interactions between neural populations (e.g., brain regions). Ideally, the recovered network can differentiate between direct and indirect interactions, and the orientation and strength of such interactions. A common strategy is to collect brain signals from a set of neural populations (termed “nodes” from here on) and compute a number of statistical association tests on the signals to infer the connectivity profile governing the set of brain nodes.

For brain signals collected using functional magnetic resonance imaging (fMRI) the most popular method to estimate associations and define networks is Pearson bivariate correlation. A significant bivariate correlation coefficient between two nodes will imply a connection or edge between those two nodes. This method is fast to compute, every scientific software has a function to do it, and straightforward statistical tests are available. Nevertheless, for the task of recovering a connectivity architecture it has important limitations. This is especially clear from a causal inference perspective, which provides a variety of well-developed concepts that are useful for illustrating the limitations with typical FC research (Pearl, 2009; Reid et al., 2019; Spirtes et al., 2000). First, if two nodes are not connected but both have a common cause (a confounder) such as A ← C → B, where C is the common cause, A and B will be correlated and a spurious or false positive edge will be defined between them (**Figure 1a**). False positive errors from common causes are costly for various reasons, for example, after observing a spurious edge a researcher could end up investing in an experiment to affect B by manipulating A, which clearly will result in no observed effect and waste time and resources.

**Figure 1.**
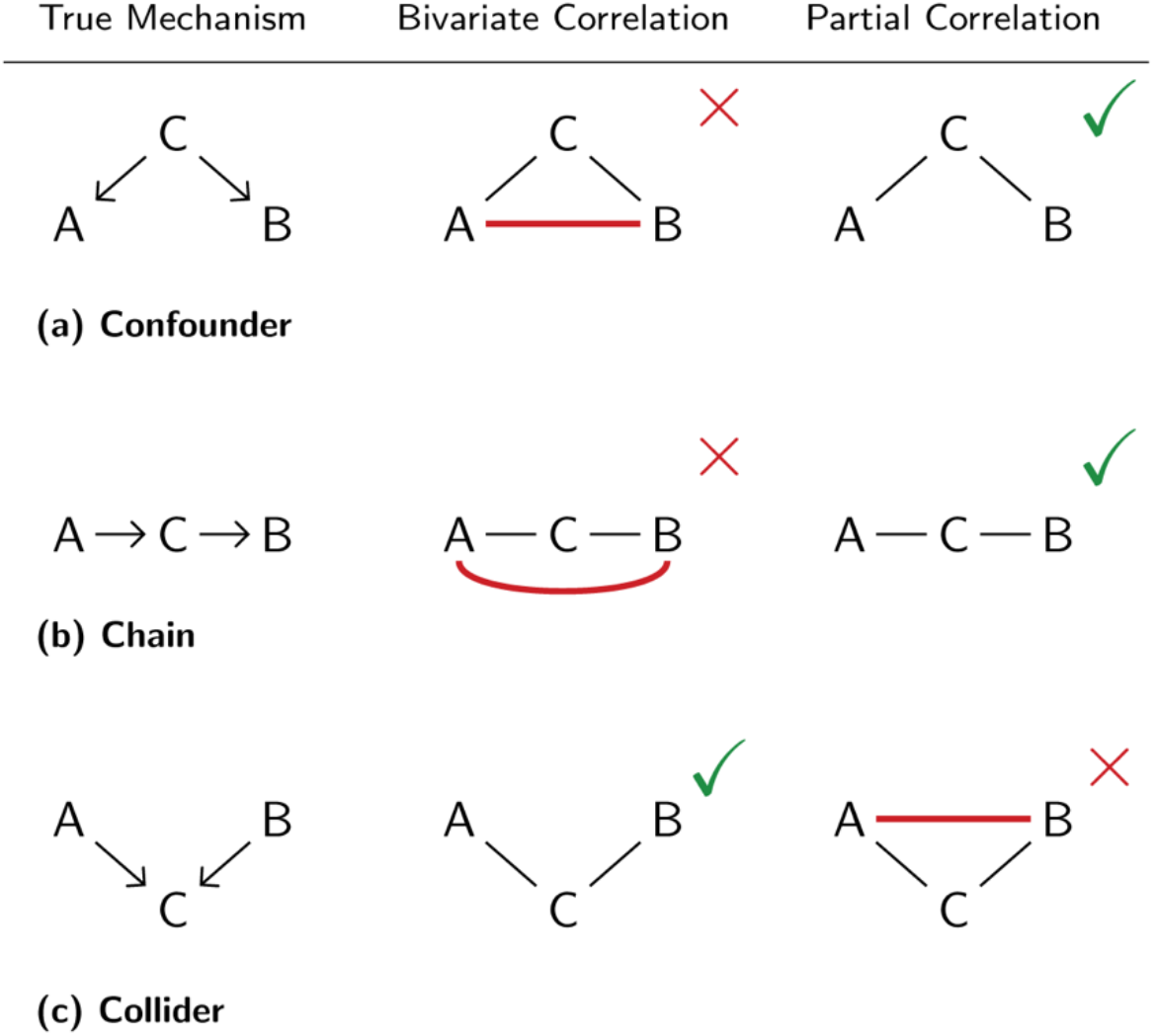
The pattern of spurious causal inferences for bivariate and partial correlations. Switching from correlation to partial correlation improves causal inference (but is not perfect). We propose integrating inferences from both correlation and partial correlation, which we predict will produce further improvements to causal inferences. Red lines indicate spurious causal inferences. Note that, in the case of a collider, when A → C and B → C are positive then the spurious A – B connection induced by partial correlation will be negative (this becomes relevant in the Results. See also Figure 6).

Second, if a bivariate correlation is observed between two nodes A and B, it is not possible to determine if this correlation was produced by a direct interaction A → B, an indirect interaction through other nodes (a chain; **Figure 1b**) such as A → C → B, or both a direct and an indirect interaction. In this sense, edges obtained with correlation are ambiguous about the direct or indirect nature of functional interactions. In contrast to the common cause case above, in the presence of a causal chain an experiment manipulating A to affect B will have the desired effect, yet information about the mechanism through which A is affecting B will be incomplete.

Partial correlation has been suggested as an alternative to correlation that alleviates the aforementioned problems (Smith et al., 2011, 2013) (**Figure 1a & 1b**). Partial correlation consists in computing the correlation between two nodes conditioning or controlling on the rest of the nodes in the dataset. Thus, partial correlation reveals correlations driven by variance shared uniquely by each pair of time series—relative to the set of included time series. Intuitively, partial correlation detects any direct association between two nodes after taking into account associations through indirect interactions or due to the presence of a common cause. For example, for a causal architecture A → C → B, the partial correlation of A and B conditioning on C is zero, indicating no direct interaction between A and B and thus no edge in the estimated connectivity network. In the case of a common cause structure A ← C → B, the partial correlation of A and B controlling for C is also zero, and no edge between A and B will be part of the estimated model. Conversely, the presence of an edge between two nodes in the estimated network will imply that the two nodes have a direct interaction.

In the presence of chains and confounders partial correlation is a preferable alternative to correlation for the goal of estimating a connectivity network, as the method is able to differentiate between direct and indirect interactions and avoid spurious edges (**Figure 1a & 1b**). However, partial correlation has an important limitation, for a causal structure A → C ← B, where C is a common effect and A and B are unrelated (i.e., C is a collider), the partial correlation of A and B conditioning on C will be non-zero and thus a spurious edge between A and B will be included in the inferred network (**Figure 1c**). Importantly, such spurious association also arises when conditioning on any other node that is an effect of a collider (Pearl, 1986). This “conditioning on a collider” effect is well-known in the causal inference and machine learning literatures (Bishop, 2006; Hernán, Hernández-Díaz, & Robins, 2004; Pearl, 1986; Spirtes et al., 2000), with application to inferring causal directionality based on the principle of testing for conditional independence (Chickering, 2002; Meek, 1995; Spirtes & Glymour, 1991).

An example may be helpful for understanding why conditioning on a collider creates a spurious association between two unrelated nodes. Assume that nodes A, B and C have two states: active or not-active, and that for node C to be active it requires both nodes A and B to be active. A researcher that only analyzes data for the states of node A and node B will reach the correct conclusion that A and B are not associated. In other words, information about the state of A does not provide any information about the state of B and vice versa. But, if the researcher also takes into account the state of C, which formally entails controlling or conditioning on C, then information about the state of A together with information about the state of C will provide information to correctly infer the state of B. For example, if A is active and C is not-active then we can correctly infer that B is not-active. This implies that region A and region B will be associated conditional on region C. The above example assumes discrete variables but the phenomenon equally holds with continuous variables.

In the presence of a collider structure A → C ← B, correlation (which does not condition on any other node) will correctly infer the absence of an edge between A and B, while partial correlation, due to conditioning on the collider C, will incorrectly infer a spurious edge between A and B (**Figure 1c**).

The behavior of correlation and partial correlation regarding confounders, colliders and chains, suggest the possibility of combining the inferences from both methods to minimize the presence of false positive edges and at the same time disambiguate between direct and indirect interactions.

We recently proposed an approach (Reid et al., 2019) to combine bivariate correlation and partial correlation to improve causal inferences, which is developed and tested for the first time here. The basic idea underlying the approach was briefly mentioned as a possibility by Smith (2012), and was inspired by existing methods that use conditional independence to infer causality (Smith, 2012; Spirtes et al., 2000). The approach involves using partial correlation to estimate an initial connectivity network, followed by checking if any of those connections has a bivariate correlation coefficient that is zero. A zero correlation will indicate the possible presence of a spurious association due to conditioning on a collider and thus the corresponding edge will be deleted from the initial network. The partial correlation step is intended to avoid spurious edges from confounders and chains, while the bivariate correlation step is meant to avoid spurious edges from conditioning on colliders. We refer to this approach as “combined functional connectivity” or combinedFC.

Properly, spurious connections arising from confounders, colliders and chains are not specific problems of bivariate correlation and partial correlation but are present for any method that tries to infer a causal mechanism from statistical associations (Spirtes et al., 2000). Measures such as mutual information and conditional mutual information, for example, will produce spurious edges in the presence of a confounder or a collider respectively. Here we focus on linear bivariate correlation and partial correlation since they are two of the most-used methods to infer brain connectivity from functional MRI data (Cole et al., 2016; Marrelec et al., 2006; Reid et al., 2019; Ryali et al., 2012). Similar considerations apply to all brain measurement methods, such as electroencephalography, magnetoencephalography, or multi-unit recording.

We implement combinedFC and compare its accuracy to bivariate correlation and partial correlation using simulations under different conditions. We then apply the three methods to empirical fMRI data from the Human Connectome Project (HCP) (Van Essen et al., 2013) to illustrate the differences between the recovered functional connectivity networks. This demonstrates that it matters in practice which method is used. For reproducibility, results are available as a Jupyter notebook at github.com/ColeLab/CombinedFC.

Here we applied combinedFC to resting-state fMRI but the method can also be used to recover connectivity networks from task-based fMRI. Formally, combinedFC is a general method that can recover functional connectivity networks as long as there are statistical conditional independence measures appropriate for the associational (linear or non-linear) and distributional (e.g., Gaussian) features of the data under consideration.

## Materials and Methods

CombinedFC builds an initial connectivity network using partial correlations to avoid spurious edges produced by confounders and causal chains, and then removes spurious edges arising from conditioning on colliders if the corresponding bivariate correlations are judged as zero. We implement the method as follows:

Partial correlations are computed using the inverse of the covariance matrix for the set of variables of interest, also known as the precision matrix P. The partial correlation coefficient for two nodes’ time series A and B conditioning on **C**, the set including all nodes in the dataset except A and B, is equal to: *r*_AB|**C**_= −P_AB_ /*sqrt*(P_AA_P_BB_), where *sqrt*( ) indicates the square root function and P_AB_ the entry for node A and B in the precision matrix. This is mathematically equivalent to computing the bivariate correlation of each pair of nodes’ time series after regressing out (controlling for) all other nodes’ time series. If the dataset has more datapoints than variables computing the precision matrix is a computationally efficient way to obtain the partial correlations for all the pairs of variables in the dataset.

To determine statistical significance, the partial correlation coefficients *r*_AB|**C**_ are transformed to the Fisher *z* statistic *Fz* = [*tanh*^−*1*^(*r*_AB|**C**_) − *tanh*^−*1*^(*r*_AB|**C**_^Ho^)]*sqrt*(N-|**C**|-3), where *r*_AB|**C**_^Ho^ is the partial correlation coefficient under the null hypothesis, N is the number of datapoints and |**C**| is the number of nodes in the conditioning set **C**. The *Fz* statistic has a distribution that approximates a standard normal with mean 0 and standard deviation 1, and is used in a two-sided *z*-test for the null hypothesis of zero partial correlation, *r*_AB|**C**_^Ho^ = 0, at a selected *α* cutoff. For *Fz*_*α*/2_, the value corresponding to the *α* cutoff in a two-sided *z*-test, if *Fz* ≥ +*Fz*_*α*/2_ or *F*z ≤ − *F*z_*α*/2_ the partial correlation is considered significantly different from zero and an edge between A and B is added to the initial network with a weight equal to *r*_AB|**C**_.

To check for spurious edges caused by conditioning on a collider in the partial correlation step, the bivariate Pearson correlation coefficient *r*_AB_ is computed for each pair of connected nodes A and B in the initial network. In contrast to the partial correlation step in which edges are added to the network, in the correlation step edges are removed if *r*_AB_ = 0. For bivariate correlation coefficients *r*_AB_, the above formula for *Fz* reduces to *Fz* = [*tanh*^−*1*^(*r*_AB_) − *tanh*^−*1*^(*r*_AB|**C**_^Ho^)]*sqrt*(N-3), since bivariate correlation does not condition on other nodes and so the size of the conditioning set |**C**| = 0. A two-sided *z*-test for the null hypothesis of *r*_AB|**C**_^Ho^ = 0 is conducted at a chosen *α* cutoff. For *Fz*_*α*/2_, the value corresponding to the *α* cutoff in a two-sided test, if *Fz* < +*F*z_*α*/2_ or *F*z > −*F*z_*α*/2_ the bivariate correlation is considered *not* significantly different from zero and the corresponding edge between A and B is removed from the initial network.

### Simulation methods

The performance of combinedFC, partial correlation and correlation is tested using data generated from linear models of the form X = WX + E, where X = {X_1_, X_2_, … , X_v_} is a vector of v variables, E = {E_1_, E_2_, … , E_v_} is a vector of v independent noise terms, and W is a matrix of connectivity coefficients with diagonal equal to zero to represent no self-loops. If the entry W_ij_ ≠ 0, then the model implies a direct causal interaction X_j_ → X_i_, otherwise X_i_ and X_j_ are not directly connected.

To simulate data for a linear model the above equation can be expressed as X = (I − W)^−1^E, where I is the identity matrix, and datapoints for the X variables are obtained by specifying the coefficients of the connectivity matrix W and datapoints for the E noise terms. A common strategy is to sample noise terms E from a standard normal distribution with mean 0 and standard deviation 1. But to simulate more realistic scenarios it is possible to implement a pseudo-empirical approach to define the E noise terms and produce synthetic variables X that better resemble the empirical data of interest. The approach consists in selecting an empirical dataset from the domain of interest and randomize the variable labels and each variable vector of datapoints individually, with the aim of destroying existing associations while keeping the marginal distributional properties of the empirical variables. The newly randomized empirical variables are used as noise terms E, and together with W define synthetic datapoints for X by the above equation. In these simulations pseudo-empirical noise terms E were built using fMRI resting state data from a pool of 100 random subjects of the HCP and parcellated into 360 brain cortex regions (Glasser et al., 2016) (see subsection below).

The simulation of the coefficient matrix W consists in the definition of a connectivity architecture (i.e., the non-zero entries in W) and the choice of coefficient values for the non-zero entries. Two different causal graphical models are used to define connectivity architectures. The first model is based on an Erdos-Renyi process (Erdős & Rényi, 1960) and produces architectures with a larger proportion of confounders than colliders. In contrast, the second model is based on a power-law process (Goh, Kahng, & Kim, 2001) and generates architectures with a larger proportion of colliders than confounders. These two models illustrate the performance of combinedFC in conditions in which partial correlation will perform better than correlation and conditions where the opposite is true.

The coefficient values for W were sampled from a uniform distribution with an interval from −1 to 1. To avoid zero or close to zero coefficients, values in the interval (−0.1, 0) were truncated to −0.1 and those in the interval [0, +0.1) were truncated to +0.1.

We analyzed the performance of the methods across simulations that vary four parameters: number of datapoints = {250, 600, **1200**}, number of regions = {50, **200**, 400}, connectivity density or percentage of total possible edges = {**5%**, 10%, 20%}, and *α* cutoff for the significance of the two-sided null hypothesis tests = {0.001, **0.01**, 0.05}. When one parameter is varied the other three are fixed to the value in bold. For example, in the simulations where the number of datapoints varies, the number of regions is fixed to 200, the connectivity density to 5%, and the *α* cutoff value to 0.01.

To compare the performance of the three methods in recovering the true connectivity architectures, we used precision and recall as measures of effectiveness (Rijsbergen, 1979).

Precision is the proportion of true positives or correctly inferred edges out of the total number of inferences: *precision* = *true positives* / [*true positives* + *false positives*]. Precision ranges from 0 to 1 and a value of 1 implies no false positives (**Figure 2b & 3b**). Recall is the proportion of correctly inferred edges out of the total number of true edges: *recall* = *true positives* / [*true positives* + *false negatives*]. Recall ranges from 0 to 1 and a value of 1 implies no false negatives (**Figure 2b & 3b**). Given that correlation and partial correlation do not recover information about the causal direction of edges, precision and recall only reflect the accuracy of the methods to recover edges regardless of their orientation. We instantiate 100 times each simulated condition and report averages and standard error bars for precision and recall across the runs.

**Figure 2.**
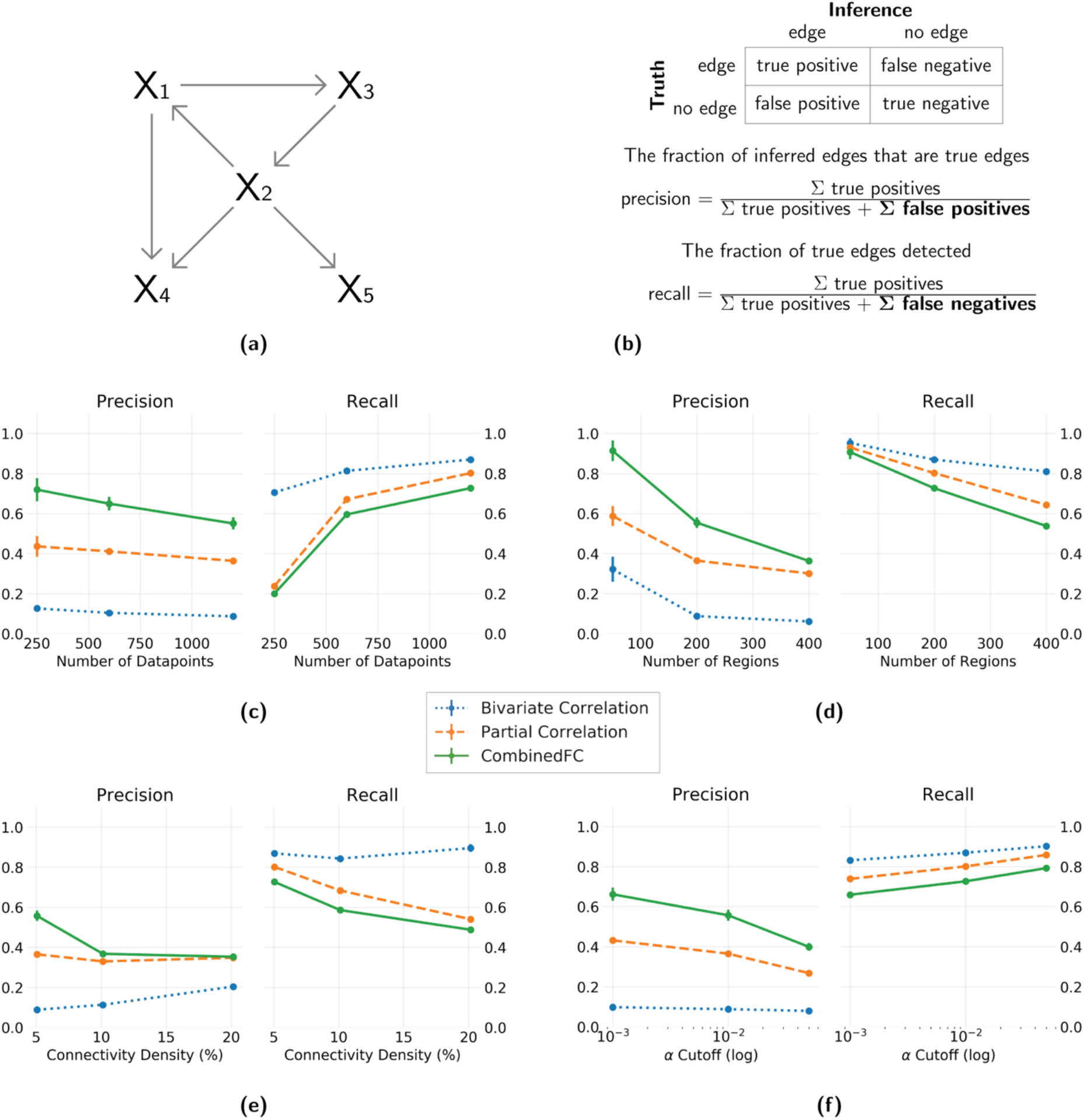
Precision and recall for simulated networks with a larger number of confounders and chains than colliders. **(a)** An example of a 5 node network generated with an Erdos-Renyi process, with more confounders and chains than colliders. **(b)** Formulas for precision and recall based on the sum of true positive, false positive and false negative inferred edges, relative to a true network. Results show average and standard deviation across 100 instantiations. Four different parameters are varied independently: **(c)** number of datapoints = {250, 600, **1200**}, **(d)** number of regions = {50, **200**, 400}, **(e)** connectivity density = {**5%**, 10%, 20%} and **(f)** *α* cutoff for the significance test = {0.001, **0.01**, 0.05}. In panel f the values are plotted in logarithmic scale for better visualization. When one parameter was varied the other three were fixed at the value in bold.

**Figure 3.**
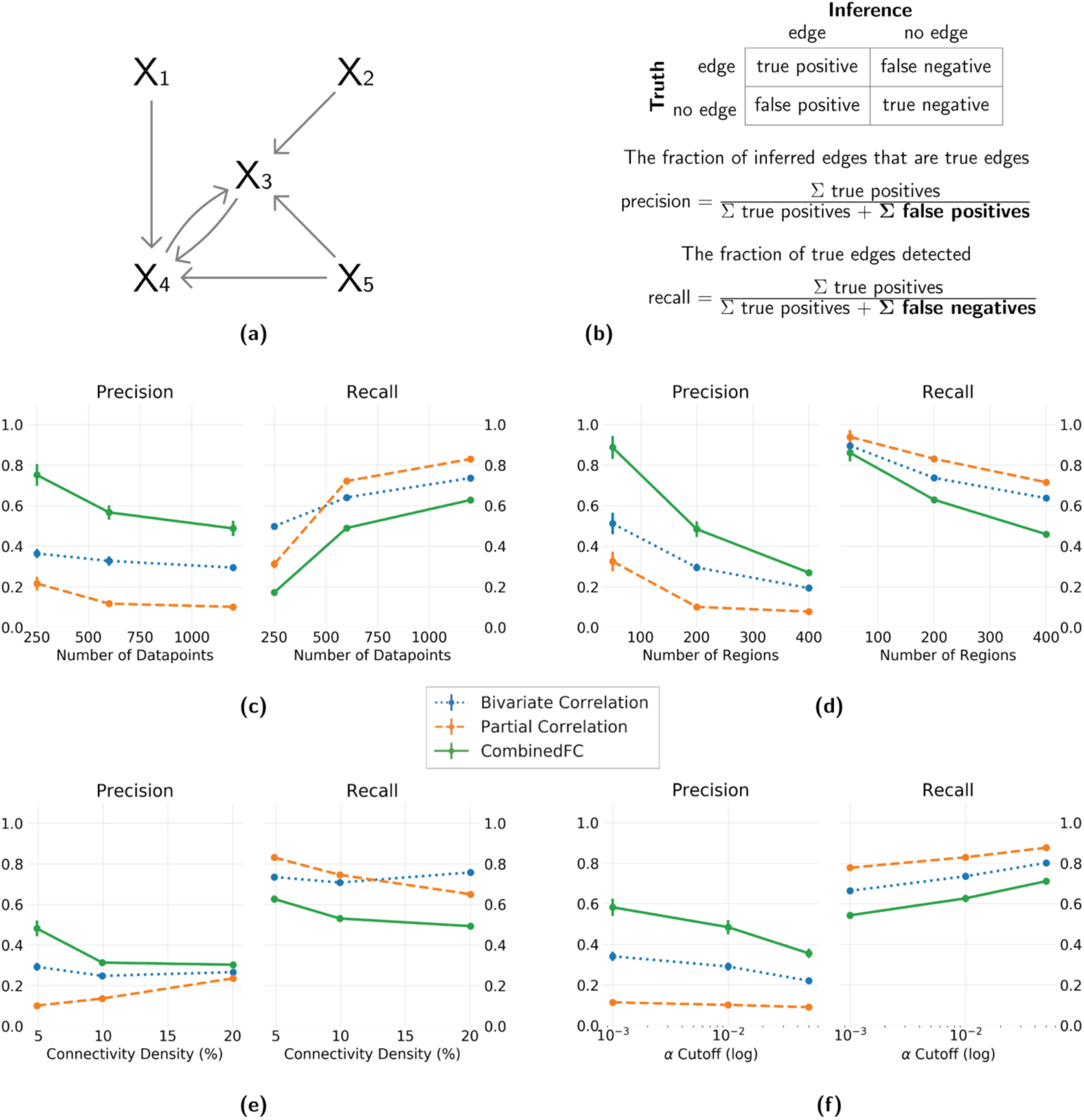
Precision and recall for simulated networks with a larger number of colliders than confounders and chains. **(a)** An example of a 5 node network generated with a power law process, with more colliders than confounders and chains. **(b)** Formulas for precision and recall based on the sum of true positive, false positive and false negative inferred edges, relative to a true network. Results show average and standard deviation across 100 instantiations. Four different parameters are varied independently: **(c)** number of datapoints = {250, 600, **1200**}, **(d)** number of regions = {50, **200**, 400}, **(e)** connectivity density = {**5%**, 10%, 20%} and **(f)** *α* cutoff for the significance test = {0.001, **0.01**, 0.05}. In panel f the values are plotted in logarithmic scale for better visualization. When one parameter was varied the other three were fixed at the value in bold.

### Empirical fMRI analysis methods

For the pseudo-empirical simulations described above and for the empirical analysis we use fMRI resting state data from a pool of 100 random subjects from the minimally preprocessed HCP 1200 release (Glasser et al., 2013), with additional preprocessing following Ito et al. (2019) and Ciric et al. (2017). HCP data was collected in a 3T Siemens Skyra with TR = 0.72 s, 72 slices and 2.0 mm isotropic voxels. For each subject 4 resting state fMRI scans were collected, each lasting 14.4 minutes, resulting in 1200 datapoints per scan. The data was parcellated into 360 brain cortex regions (180 per hemisphere) defined in Glasser et al. (2016). Additional preprocessing on the parcellated data include removing the first 5 datapoints of each scan, demeaning and detrending the time series and performing nuisance regression with 64 regressors to remove the confounding effect of various motion and physiological artifacts. No global signal regression was implemented. Finally, the time series of the 360 regions were individually standardized to bring them to the same scale with mean 0 and standard deviation 1. Specific details about the nuisance regression are in Ito et al. (2019).

For the empirical fMRI data, we illustrate the use of combinedFC in a group level analysis and compared its result to bivariate correlation and partial correlation. We only use the first resting state session data (1195 datapoints x 360 regions) for each subject analyzed. The goal of the group analysis is to obtain a connectivity network reflecting group average significant connections. The group analysis for bivariate correlation and partial correlations follows Smith et al. (2013). Let M^s^ for s = {1, … , n} denote a group of n subjects bivariate correlation (or partial correlation) matrices, and M^g^ the group average connectivity matrix we want to infer. First, each M^s^ is transformed into a Fisher *z* statistics matrix F^s^. Then, for each ij entry we compute the group average 1/n Σ_s_(F^s^_ij_) and perform a two-sided one-sample *t*-test for the null hypothesis H_0_ : 1/n Σ_s_(F^s^_ij_) = 0. Finally, if H_0_ is rejected at the chosen *α* value, the ij entry of the group connectivity matrix M^g^ is defined as M^g^_ij_ = 1/n Σ_s_(M^s^_ij_), otherwise M^g^_ij_ = 0.

For combinedFC, the group analysis starts by computing an initial partial correlation group average connectivity matrix M^g^ as above. Then, for each non-zero entry M^g^_ij_ ≠ 0 we perform a two-sided one-sample *t*-test for the null hypothesis H_0_ : 1/n Σ_s_(ϕ^s^_ij_) = 0, where ϕ^s^_ij_ is the Fisher *z* transform of the bivariate correlation between node i and node j for subject s. In other words, we want to determine if the group average bivariate correlation between two nodes is significantly different from zero or not. As mentioned before, the correlation step looks to remove connections with non-zero partial correlation but zero bivariate correlation since that indicates a possible spurious connection from conditioning on a collider. So, if the H_0_ is *not* rejected at the chosen *α* value—meaning that the group average correlation is not significantly different from zero—we remove the connection by setting M^g^_ij_ = 0, otherwise we keep the initial value of M^g^_ij_.

An alternative approach for inferring if bivariate correlations are zero (or close enough to zero that we consider them to be null effects) is to use an equivalence test (Goertzen & Cribbie, 2010; Lakens, 2017). Formally, it is not appropriate to use non-significance to infer that the null effect was true. This is clear in the case of high uncertainty (e.g., high inter-subject variance), since even large correlations would be considered non-significant. Non-significance implies that we do not have sufficient evidence of a correlation, not that we have strong evidence that the correlation is zero. This is also clear in the case of very small effects with large sample size, in which extremely small correlations are still considered significantly non-zero. Significance in this case implies that we have sufficient evidence of a correlation, but that correlation might be so small that it is not above zero in a meaningful sense. For instance, *r*_AB_ = >0.20 is such a small effect size that only 4% of linear variance is shared between time series. Unlike non-significance in a two-sided one-sample *t*-test, equivalence tests allow one to properly infer that the null effect is true by choosing a minimum effect size of interest (e.g., we consider *r*_AB_ to be zero, if *abs*(*r*_AB_) < 0.20, where *abs*( ) is the absolute value function). Equivalence tests are very straightforward, simply using standard null hypothesis testing (e.g., a *t*-test) to determine if an effect is significantly closer to zero than the chosen minimum effect size of interest. We implement the equivalence test in the group analysis as follows:

In an equivalence test to determine if a group average bivariate correlation is zero, two one-sided one-sample *t*-tests are conducted. First, *Fz*^L^ and *Fz*^U^ are defined as the Fisher *z*-transformed values of the negative (lower bound) and positive (upper bound) of a chosen minimum bivariate correlation coefficient of interest—minimum effect of interest. The lower bound *t*-test is a right-sided test for the null hypothesis H_0_^L^ : 1/n Σ_s_(F^s^_ij_) = *Fz*^L^, and alternative hypothesis H_A_^L^ : 1/n Σ_s_(F^s^_ij_) > *Fz*^L^. The upper bound *t*-test is a left-sided tests for the null hypothesis H_0_^L^ : 1/n Σ_s_(F^s^_ij_) = *Fz*^U^, and alternative hypothesis H_A_^L^ : 1/n Σ_s_(F^s^_ij_) < *Fz*^U^. For a selected *α* cutoff, if both H_0_^L^ *and* H_0_^U^ are rejected, the equivalence test concludes that significantly *Fz*^L^ < 1/n Σ_s_(F^s^_ij_) < *Fz*^U^. This result implies that the group average bivariate correlation of node i and node j is inside the bounds of the minimum effect of interest and will be judged as zero. In combinedFC this result implies setting M^g^_ij_ = 0, otherwise keeping the initial value of M^g^_ij_.

Code to implement combinedFC at the individual and group level is available as a toolbox at github.com/ColeLab/CombinedFC.

## Results

### Validating combinedFC using simulations

As combinedFC benefits from the capacity of partial correlation to avoid false positive edges from confounders and chains, and from the capacity of correlation to avoid false positive edges from conditioning on colliders, it is expected to have a lower number of false positives than either of the two methods alone. Consistent with this the results of all simulations show that combinedFC has better precision than both bivariate and partial correlation. As combinedFC starts with the partial correlation connectivity network and does not add any more edges, its number of true positives is the same as for partial correlation, so any improvement in precision relative to partial correlation necessarily comes from a reduction in false positives. As expected, partial correlation has a better precision than bivariate correlation in the simulations from models with a large number of confounders and chains (**Figure 2**), while bivariate correlation precision is higher in simulations from models with a large number of colliders (**Figure 3**). CombinedFC precision is the highest for both types of graphical models.

CombinedFC’s recall upper bound is determined by partial correlation’s recall. CombinedFC true positives are the same as partial correlation true positives, so any reduction in recall relative to partial correlation comes from an increase in the number of false negatives. An increase in false negatives means that combinedFC is incorrectly removing some true edges.

Since combinedFC is based on partial correlation and bivariate correlation, increasing datapoints (**Figure 2c & 3c**) will also have a positive impact on precision and recall. We found that combinedFC precision and (to a larger degree) recall were improved by increasing datapoints, likely due to more statistical power to correctly detect the presence of true edges.

CombinedFC has a strong positive effect in precision for all the number of regions we simulated (**Figure 2d & 3d**). This effect diminishes to a degree in the larger model we tested, possibly due to the fact that in larger simulated networks there is a higher probability that nodes with common causes, common effects or indirect interactions are also directly connected, such that combinedFC removes fewer false connections.

**Figure 2e & 3e** show an excellent precision improvement of combinedFC in sparse problems (low connectivity density). This is expected since in sparse causal architectures any two nodes have a higher probability of not being connected, while still potentially having a common cause, common effect or an indirect interaction (such that combinedFC removes more false connections). As the true causal architecture becomes denser the benefit in precision from combinedFC is reduced, since now—as with the simulations with large number of regions— there is a higher probability that nodes with common causes, common effects or indirect interactions are also directly connected (such that combinedFC removes fewer false connections).

Changes in recall and precision of combinedFC can be achieved by changing the *α* cutoff for the significance tests. **Figure 2f & 3f** show that in these simulations a smaller *α* value improves the precision and decreases the recall for combinedFC. In the opposite direction, by choosing a larger *α* the hypothesis tests will be more lenient by increasing the recall (more true edges will be judged significant) at the cost of a lower precision (more false edges will be judged significant). When using combinedFC the preference between precision or recall is a decision that depends on the researchers’ goals. Scientists generally tend to value precision over recall, since false positives are thought to be more costly to scientific progress than false negatives (Parascandola, 2010).

For the network inference problem, simple and multiple linear regression can be used to obtain measures of unconditional and conditional linear association for a node and a set of node regressors. This makes them straightforward alternatives to partial correlation and bivariate correlation to implement combinedFC. In this case, we would first compute the multiple regression of each node on the rest of the nodes, for example X_1_ = β_0_ + β_2_X_2_ + … + β_v_X_v_ + E_1_. Then, for each non-zero multiple regression coefficient β_j_ ≠ 0 we would compute the *simple* regression between the node and the corresponding regressor X_1_ = γ_j_X_j_ + E_1_. The simple regression coefficient γ_j_ is a measure of unconditional association and thus an alternative to the bivariate correlation coefficient for the collider check. If two nodes have a non-zero multiple regression coefficient β_j_ ≠ 0, but a zero simple regression coefficient γ_j_ = 0, then we have evidence of a spurious edge from conditioning on a collider. We applied a combinedFC implementation with ordinary least squares multiple linear regression and simple linear regression to our simulations, and obtained equivalent inference precision and recall to the ones of combinedFC with bivariate correlation and partial correlation. This is expected given the theoretical relationship between the methods. We include this linear regression implementation of combinedFC in the accompanying toolbox.

### Applying combinedFC to empirical resting state fMRI data

We used the same pool of 100 HCP subjects from the pseudo-empirical simulations to perform group analyses and assess whether it mattered in practice which method (bivariate correlation, partial correlation, or combinedFC) is used in empirical fMRI data analyses. This also allowed us to assess the relative performance of the methods as a function of sample size, and as a function of how false connections are identified.

We began by measuring the sensitivity of the three approaches to increasing number of subjects in the group. Groups with 10, 40, 70 and 100 subjects were evaluated. For each case the three functional connectivity methods were applied as described in Materials and Methods. A significance cutoff of *α* = 0.01 was used for the significance tests of correlation, partial correlation and combinedFC, for all group sizes.

**Figure 4** shows the number of edges inferred by each method (bivariate correlation excluded from visualization) at each of the four group sizes analyzed. The number of inferred edges for all methods increase with the size of the group. The results for bivariate correlation are not plotted in **Figure 4**, since they are one order of magnitude larger than for the other methods, but they are included here for completeness: 45,120 edges (with 10 subjects); 58,019 (40); 60,442 (70); 61,206 (100). The results labeled as “CombinedFC non-significance”, refer to combinedFC using non-significant correlations for the collider check as described in Materials and Methods. As can be seen, the number of inferred edges for this implementation of combinedFC are very similar to the ones from partial correlation. One reason behind this result is that in group analyses with high statistical power, very small spurious bivariate correlations (e.g., *r*_AB_ = 0.10) may be judged significant and thus combinedFC will not judge them as spurious edges from conditioning on a collider.

**Figure 4.**
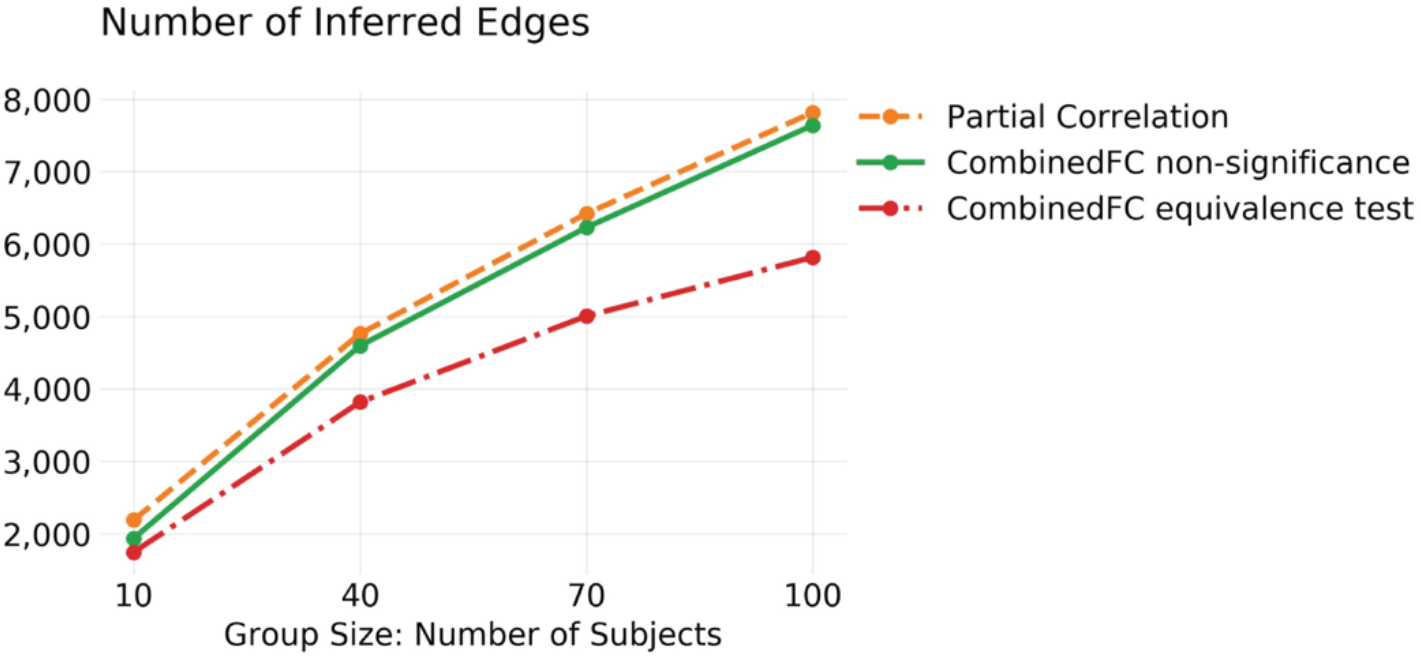
Empirical data analyss comparing strategies to identify false connections due to colliders. Number of inferred group edges for partial correlation and combinedFC implemented with non-significant correlations and with equivalence tests for the collider check. Only the equivalence test strategy identifies more false connections with a larger sample size, consistent with more data providing greater evidence of no (or a very small) bivariate correlation in these cases. See text for results with bivariate correlation.

As mentioned in Materials and Methods, an alternative approach for inferring zero correlations is to use an equivalence test. By choosing a minimum effect of interest, an equivalence test allows us to make a significance judgment of zero correlation and overcome the problem of very small significant correlations described above for the non-significance implementation. We applied combinedFC with an equivalence test, choosing a minimum bivariate correlation coefficient of interest of 0.2 and a significance cutoff of *α* = 0.01. Results are shown in **Figure 4** as “CombinedFC equivalence test”. CombinedFC with an equivalence test inferred a smaller amount of edges than combinedFC with the non-significance judgement. As the group size increases, the equivalence tests gain more statistical power to correctly judge the presence of zero mean group effects (Lakens, 2017) and thus combinedFC becomes more effective in removing potential spurious edges from conditioning on colliders.

For illustration and comparison we plotted in **Figure 5** the connectivity networks for the 100 subjects group analysis. Connection weights represent mean values across 100 subjects. The network for combinedFC with non-significance judgments is not plotted since, as explained above and shown in **Figure 4**, the results are very similar to partial correlation. The rows and columns of the matrices are ordered according to 12 functional networks described in Ji et al., (2019).

**Figure 5.**
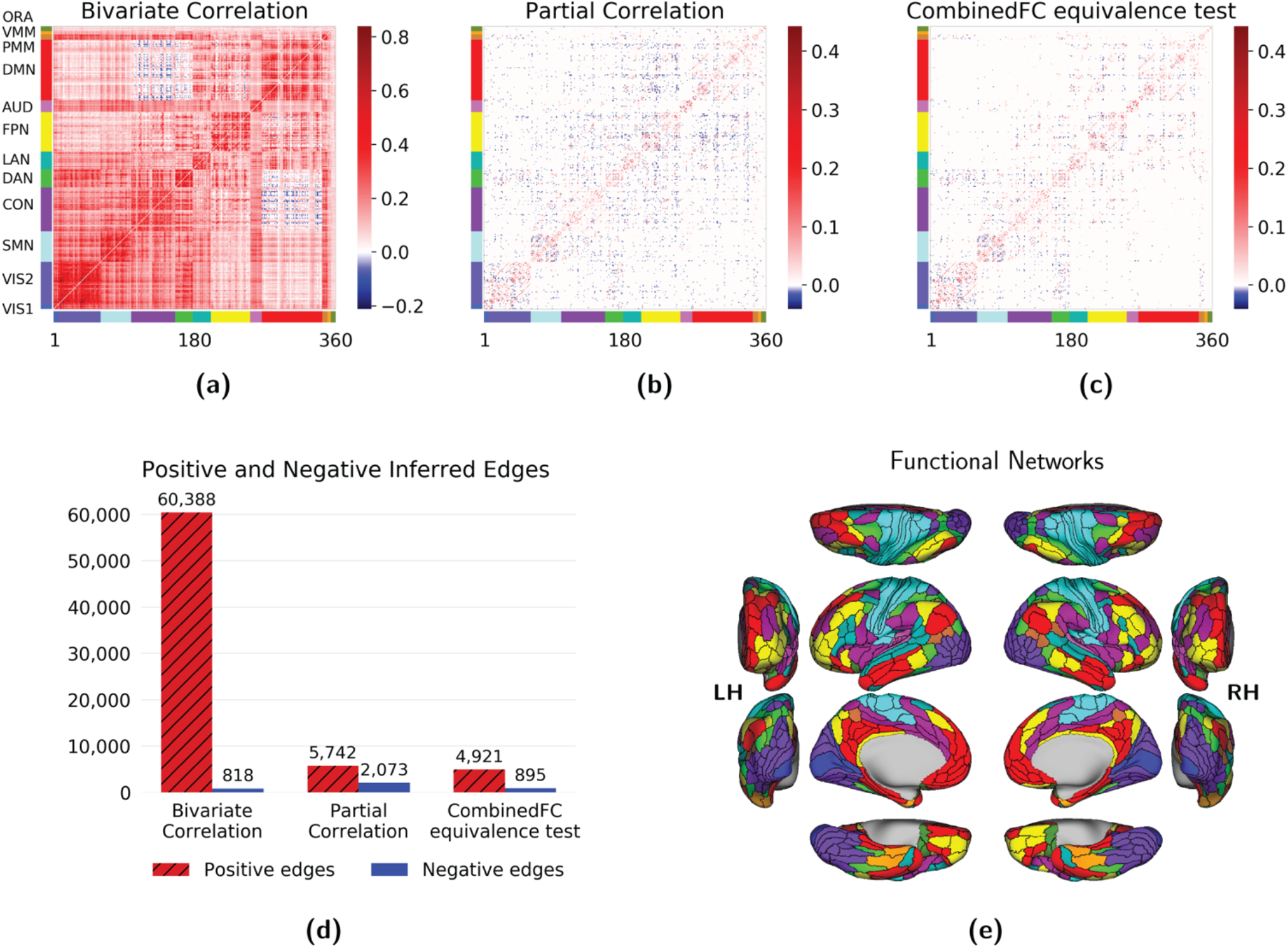
Comparison of bivariate and partial correlation with combinedFC using empirical resting state fMRI data. Results for **(a)** bivariate correlation, **(b)** partial correlation and **(c)** combinedFC with equivalence tests. **(d)** Number of positive and negative inferred edges by each method. Partial correlation removed a large number of positive edges and increased the number of negative edges relative to correlation. This increase may come from spurious edges from conditioning on colliders with same sign associations. Consistent with the removal of spurious partial connections caused by colliders, combinedFC removed 57% of the negative partial correlations and ended up with a number of negative edges close to the one of the bivariate correlation matrix, for which no spurious edges from colliders are present. **(e)** The 360 regions of interest are ordered according to 12 functional networks defined in Ji et al., (2019) using bivariate correlation: VIS1: primary visual; VIS2: secondary visual; SMN: somatomotor; CON: cingulo-opercular; DAN: dorsal attention; LAN: language; FPN: frontoparietal; AUD: auditory; DMN: default mode; PMM: posterior multimodal; VMM: ventral multimodal; ORA: orbito-affective.

In the 100 subjects group analysis, correlation (**Figure 5a**) produced a dense network with 61,206 significant edges (out of 64,620 possible), while partial correlation (**Figure 5b**) inferred a sparser model with 7,815 significant edges (**Figure 5e**). This massive reduction in inferred edges from using partial correlation likely reflects the widespread presence of confounders and causal chains among brain regions. Partial correlation removes false edges from confounders and indirect connections, producing a brain connectivity network in which edges between regions can be interpreted (under certain assumptions) as direct connections.

It is worth noticing that the number of negative edges in the partial correlation matrix increased to 2,073 from 818 in the correlation matrix (**Figure 5d**). Two unconnected nodes will have a negative spurious partial correlation from conditioning on a collider if their connectivity coefficients with the collider have the same sign. In contrast, they will have a positive spurious partial correlation if their associations have opposite signs (Reid et al., 2019; Smith, 2012) (**Figure 6**). Anatomical studies in non-human primates have established that most long-range cortico-cortical connections are positive (i.e., glutamatergic) (Barbas, 2015), such that we can reasonably assume that most true connections among brain regions have the same sign (positive). This suggests that—assuming fMRI data properly reflects underlying neural signals (Lee et al., 2010)—most spurious partial correlations will be negative. Consistent with the removal of spurious partial connections caused by colliders, combinedFC (**Figure 5c**) reduced to 895 the number of negative edges from the initial 2,073 in the partial correlation matrix (**Figure 5e**). It is interesting that the final number of negative edges in the combinedFC matrix is close to the number of negative edges in the bivariate correlation matrix, in which no spurious edges from conditioning on collider are present.

**Figure 6.**
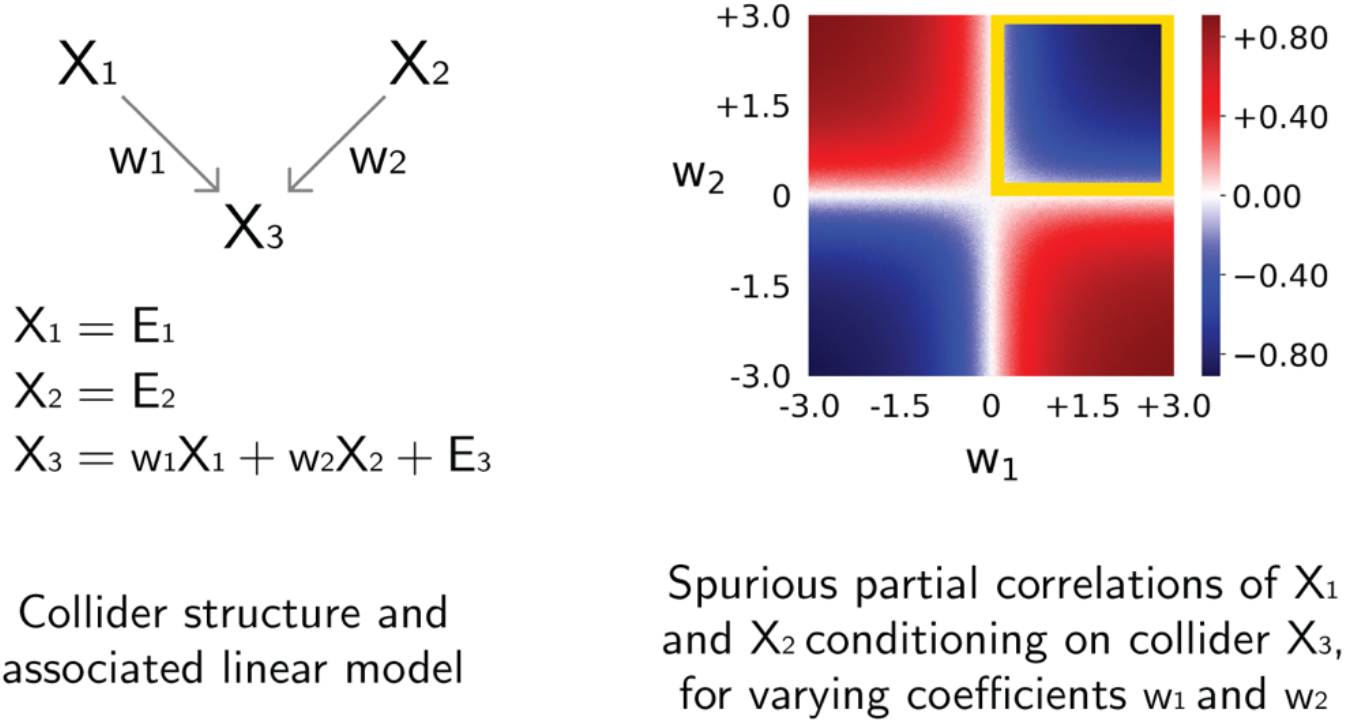
Positive or negative spurious partial correlations when conditioning on a collider. The left panel shows a collider causal structure from regions X_1_ and X_2_ to X_3_ and an associated linear model with connectivity coefficients w1 and w2, and Gaussian noise terms E. We simulated data from this linear model for values of w1 and w2 from −3 to +3, and report in the right panel the resulting spurious partial correlations of X_1_ and X_2_ conditioning on X_3_ for combinations of values of w1 (x-axis) and w2 (y-axis). When both connectivity coefficients have positive values the resulting spurious partial correlation will be negative (blue region indicated by a yellow square). Negative spurious partial correlations are also observed if both connectivity coefficients are negative. In contrast, when the connectivity coefficients have opposite sign, the resulting spurious partial correlation will be positive (red regions).

The group analysis reported in **Figure 5** was repeated for the same set of 100 subjects but using the second session of resting-state fMRI available in the HCP dataset. The results for the second session show a similar pattern with 62,929 positive and 183 negative edges in the correlation matrix (compared to 60,388(+) and 818(−) edges in the first session); 5,673 positive and 2,072 negative edges in the partial correlation matrix (5,742(+) and 2,073(−) in the first session); and 4,951 positive and 1,005 negative edges in the combinedFC matrix (4,921(+) and 895(−) edges in the first session). These results show that partial correlation and combinedFC recovered a more stable group connectivity matrix across sessions in terms of number of positive and negative edges, while the correlation matrices differed by a larger amount across sessions, in both the number of positive and negative edges.

All together, these findings confirm the benefit—relative to either bivariate or partial correlation alone—of the combinedFC strategy to detect direct connections between regions.

## Discussion

We have shown that combinedFC provides a strategy to accurately recover connectivity networks by taking into account the way that causal relationships such as confounders, causal chains and colliders may produce spurious edges when correlation and partial correlations are used separately. Using a series of simulations varying the number of datapoints, number of regions, connection density and significance cutoff, we showed that combinedFC consistently improves the inference precision (reducing false positive edges) without considerable loss in recall (increasing false negative edges). We also presented simulations of graphical models with a majority of confounders, for which partial correlation performs better than bivariate correlation, and models with a majority of colliders, for which bivariate correlation performs better than partial correlation, and show the superior precision of combinedFC in both types of models. This result shows that partial correlation by itself is not a better strategy than correlation under all possible scenarios, and that the behavior of correlation and partial correlation used separately depend on the particular causal mechanisms governing the true network. In contrast, combinedFC takes into account the strengths and limitations of both methods and achieves a better performance regardless of the underlying causal topography of the network.

The superior precision of combinedFC in these simulations confirms the benefit of adopting a causal perspective about the data generating system of interest. In combinedFC we assume that brain signals come from a causal system that can be modelled in terms of direct causal connections between nodes, and that those direct connections give rise to patterns in the shape of confounders (A ← C → B), causal chains (A → C → B) and colliders (A → C ← B), which when incorrectly modeled can give rise to spurious edges. These causal assumptions lead to increased confidence that the combinedFC inferences are not reflecting indirect connections (because we condition for causal chains using partial correlation) nor spurious non-existent edges (because we condition for confounders using partial correlation, and account for conditioning on colliders by doing a bivariate correlation check).

Critically, because bivariate correlation is by far the most popular FC measure in fMRI research, combinedFC should be primarily assessed relative to bivariate correlation. Since the limitations with combinedFC are substantially fewer than those of bivariate correlation, widespread adoption of combinedFC would meaningfully benefit the field of FC research. To illustrate this improvement, consider the empirical fMRI application presented here. It is a whole-cortex 360 region problem for which we are trying to infer a network containing relevant information about the strength and directness of its connections. It is also a densely connected system with a high probability for the presence of confounders (e.g., a primary visual region sending information to various secondary visual regions upwards in the visual stream), causal chains (e.g., intermediate regions serving as relays in information paths from primary sensory regions to decision-making centers) and colliders (e.g., hubs that consolidate information coming from different sensory regions). The problem with using bivariate correlation for this problem is that it is impossible to disambiguate if the inferred edges represent real direct connections, indirect connections of different degrees or spurious confounded connections. In this sense, the only conclusion we can make about an edge between A and B in a bivariate correlation network is that time series A and B are associated to a certain degree without knowing anything about the mechanism producing their association. Even if we threshold a correlation matrix by correlation strength, the remaining edges cannot be disambiguated: It is possible that due to strong connections with the intermediate nodes in a chain (or with the common cause), an indirect connection (or spurious edge) results in a very strong correlation coefficient that survives the threshold. Due to the inherent causal ambiguities of bivariate correlation, we can only conclude that the nodes interact (directly or indirectly) and/or are similarly influenced by common node(s) (Reid et al., 2019).

Without causal assumptions it is not possible to overcome the ambiguities of bivariate correlation and its limitations as an informative FC method. In contrast, combinedFC uses two simple causal assumptions. The first is that by conditioning on the proper nodes we can disambiguate between direct, indirect and spurious connections. Properly, by using full partial correlation as a first step we attempt to 1) condition on intermediate nodes from causal chains to avoid edges that represent indirect connections, and 2) condition on confounders to avoid edges that represent spurious connections. The second causal assumption is that conditioning on a collider will associate two nodes that were previously independent. This assumption suggests the rule that if we found a partial correlation between two nodes but no evidence of bivariate correlation—where no conditioning is made—we will be in the presence of a spurious association and the corresponding edge should be deleted. These causal assumptions are what provide relevant information about strength and directness of associations, allowing the interpretation of edges in an inferred connectivity matrix as direct connections between nodes, thus making combinedFC a method more appropriate than bivariate correlation for the goals of FC research (Reid et al., 2019).

CombinedFC can be described as a method that builds an initial connectivity network by computing the conditional associations between each pair of nodes given the rest, to avoid spurious edges from confounders and causal chains, and then removes spurious edges arising from conditioning on colliders if the corresponding nodes are not unconditionally associated. This general description implies that we can use different methods to compute associations and conditional associations, depending on the properties of the data or other theoretical and computational considerations. The benefits of combinedFC depend on its causal assumptions and not on any particular implementation. So, as with correlation and partial correlation, combinedFC should be a better approach than any of the chosen statistical association (e.g., mutual information) and conditional association (e.g., conditional mutual information) methods used alone.

As mentioned in Results, multiple and simple regression are straightforward alternatives to partial correlation and correlation to implement combinedFC. The β_j_ coefficients of a *multiple* regression are a measure of the conditional associations between a node and each of its individual j regressors controlling for the rest. A multiple regression coefficient β_j_ = 0 will indicate that the node and its regressor j are conditionally independent given the rest of the nodes (assuming linear relationships). Conversely, β_j_ ≠ 0 will indicate a conditional association. This property of multiple regression makes it a valid alternative to partial correlation in combinedFC. In the same way, the simple regression coefficient is a measure of unconditional association and thus an alternative to bivariate correlation for the combinedFC collider check. Multiple regression has been successfully used to build resting state connectivity models from which predictions about task activations are made (Cole, et al., 2016; Ito et al., 2017) and thus it is expected that using combinedFC to remove spurious connections from these models will allow even better predictions—or at least predictions that are more causally accurate.

The general description of combinedFC does not make any assumption about the distribution of the data, temporal properties or linear relationship between the nodes. This suggests the adaptability of the strategy to different data assumptions. Next we present some future research scenarios and how combinedFC can be adapted to them.

To get reliable estimates, partial correlation requires more datapoints than nodes. The simulations and empirical data presented here satisfy this requirement. For problems where the number of nodes is considerably larger to the number of datapoints, also known as high-dimensional problems, it is necessary to apply specially-tailored methods to assess conditional associations (Bühlmann & Van De Geer, 2011), otherwise the variance of the estimators increase to infinity, making them unusable (James, Witten, Hastie, & Tibshirani, 2013). This is a common situation with fMRI, for instance, because of the typically larger number of voxels relative to datapoints.

Two of the most popular high-dimensional methods to recover networks are forms of regression regularization such as lasso (Tibshirani, 1996) and ridge regression (Hoerl & Kennard, 1970), which compute regressions with an extra regularization parameter that shrinks coefficients to zero or close to zero. Other high-dimensional alternatives are methods that estimate a regularized inverse covariance matrix from which partial correlation coefficients can be derived. Glasso (Friedman, Hastie, & Tibshirani, 2008) is possibly the most popular of these methods. Hinne, et al., (2015) introduce a Bayesian solution using priors, and BigQuic (Hsieh, Sustik, Dhillon, Ravikumar, & Poldrack, 2013) is a recent algorithm that can scale up to a million variables and has been applied to a whole-cortex voxel level problem. One more alternative for high-dimensional problems is to use dimension reduction methods (James, et al., 2013) such as principal components regression PCR (Hotelling, 1957; Kendall, 1957), in which principal components analysis (PCA) is used to obtain a low-dimensional set of components which are then used as regressors in a multiple linear regression. In high-dimensional problems the combinedFC strategy of computing conditional associations followed by simple associations is still valid, but it has to be implemented with regularization methods. For example, the first step can be computed with glasso to determine pairs of nodes that are conditionally associated, and the second step can be computed with bivariate correlation or simple regression to detect possible spurious edges from conditioning on a collider. We include a glasso implementation for combinedFC in the accompanying toolbox.

The simulations used here assume nodes interact in a linear fashion, such as X = *b*Y + E, where *b* is the association coefficient. Bivariate correlation and partial correlation reliability is guaranteed for linear problems, but if the assumption of linearity is not valid, for example X = *b*Y^2^ + E, it will be necessary to adopt non-linear association and conditional association methods. Importantly, the logic behind combinedFC is valid (in principle) for non-linear interactions, such that measures of association other than bivariate and partial correlation could be used. The conditional mutual information and mutual information test from Cover & Thomas (2012), the kernel-based conditional independence test from Zhang, Peters, Janzing, & Schölkopf (2011) and the scalable conditional independence test from Ramsey (2014), are alternatives to implement combinedFC in the presence of non-linear interactions.

Bivariate correlation and partial correlation, as used here, do not exploit temporal lag properties of brain signals. We could do this, for example, by considering a dynamic linear model X_t_ = WX_t-k_ + E_t_, where the variables are time indexed, there is a temporal lag k ≥ 1, and the W matrix encodes the temporal interactions between the variables. To make inferences about the temporal associations and conditional associations between nodes when taking temporal lags into consideration, we require methods that include assumptions about the dynamics governing the causal mechanisms. The challenge of combinedFC in the temporal domain is then to properly model the temporal dynamics of common causes, causal chains and colliders. Popular approaches model the connectivity mechanisms as structural vector autoregressive processes and try to learn a dynamic network using temporal conditional association facts (Malinsky & Spirtes, 2018; Moneta et al., 2011; Runge, 2018). Alternative methods to recover dynamical networks using different causal assumptions also have shown good results under certain conditions and some of them have been applied to neural data (Gates & Molenaar, 2012; Runge et al., 2019; Weichwald et al., 2016); and a strategy that also combines bivariate and partial temporal associations to improve causal inferences is described in Stramaglia et al., (2014).

The problems described above show the flexibility of combinedFC to different data scenarios. This flexibility derives from the fact that the benefits of combinedFC are based on its causal assumptions and not on the particular statistical association methods used to implement it. Nevertheless, there are limitations of combinedFC that arise from the presence of particular causal patterns in the true networks.

The main limitation of combinedFC is that it is not guaranteed to avoid *all* possible spurious edges from confounders and colliders. Consider a model where A and B are not directly connected but have a common cause A ← C → B, together with a common effect, A → D ← B. In the first step of combinedFC, the conditional association of A and B conditioning on C and D will be non-zero because we conditioned on the collider D, but in the second step the unconditional association of A and B will also be non-zero because of the presence of the confounder C. In this model, combinedFC will always infer a spurious edge between A and B. Notice that bivariate correlation and partial correlation used alone will also infer such spurious edge. This kind of causal pattern requires a different strategy, such as choosing conditioning sets of different sizes in an iterative way to remove spurious edges from an initially fully connected model. The Peter Clark (PC) algorithm—one of the conditional independence strategies for making causal inferences (Mumford & Ramsey, 2014; Spirtes & Zhang, 2016)— and its modifications pioneered this strategy to recover an undirected network from which logical inferences about the orientation of the edges are then made (Colombo & Maathuis, 2014; Spirtes et al., 2000). Another strategy is to use, if available, theory-based expert knowledge to define appropriate conditioning sets as subsets of the available variables, or in a more data-driven way define conditioning sets by selecting the variables more statistically associated with the pair of variables for which the conditional independence is being evaluated. The effectiveness of this strategy has been explored in Marinazzo et al., (2012) and Stramaglia et al., (2014). We recommend the PC algorithm and related approaches when it is important to avoid such cases. Notably, PC and related approaches are more complex than combinedFC, such that combinedFC might be generally preferred due to the ability for researchers to more easily understand how it works.

Cyclic interactions are other causal patterns that pose challenges to the combinedFC strategy. Assume a model where two nodes A and B are not directly connected but each one has a feedback cyclic interaction with C, such as A ⇄ C ⇄ B. Here, the node C acts both as a confounder and as a collider. In data sampled from this network, the conditional association of A and B controlling for C will infer a spurious edge due to conditioning on the collider C, and the unconditional association will also infer a spurious edge due to the confounder C. CombinedFC will incorrectly infer a spurious edge between A and B in this cyclic network. Likewise, bivariate correlation and partial correlation used alone will produce a spurious edge in this case. Making causal inferences in cyclic scenarios is an active area of research and network learning algorithms are available for particular assumptions of domain, linearity, distribution and temporal properties of the data (Runge, 2018; Sanchez-Romero et al., 2018).

There are connectivity patterns for which combinedFC will correctly infer the presence of an edge between two nodes but will incorrectly estimate the strength of the direct association. Consider three nodes A, B and C, for which combinedFC first inferred a non-zero partial correlation between nodes A and B controlling for C, and then inferred a non-zero correlation between nodes A and B. According to the combinedFC rules, these two results imply that A and B are directly connected with a strength equal to the partial correlation coefficient *r*_AB|C_. The problem is that these results are underdetermined and can be produced by three different causal structures: If the true structure is A → B with a confounder A ← C → B, then the strength of the direct association between A and B will be correctly captured by the partial correlation coefficient between A and B controlling for C, *r*_AB|C_; the same happens in the case of a chain A → C → B. But if the true structure is A → B with a collider A → C ← B, then the strength of the direct association between A and B will not be correctly captured by the partial correlation coefficient but by the bivariate correlation coefficient between A and B, *r*_AB_. In a problem like this, combinedFC will correctly infer the presence of an edge between A and B but will not be able to disambiguate the correct strength of their direct association. The ambiguity of the association strength can be resolved if information about the orientation of the edges is obtained via expert knowledge or causal learning methods (Mumford & Ramsey, 2014; Sanchez-Romero et al., 2018). For example, if the learned model from the data is A → B and A ← C → B, then a regression of node B on its two direct causes A and C will give a correct estimate of the direct association strength between B and A. As another example, if the inferred model is A → B and A → C ← B, then the regression of B onto its only direct cause A will give a correct estimate of the direct association between B and A.

Finally, as with any statistical method that tries to recover a connectivity network from conditional associations, the results of combinedFC are dependent on the variable set considered. This implies that if we remove or include new variables in the dataset, inferences about the presence or absence of previously inferred edges may change. For example, a previously inferred edge between brain regions A and B may be removed when we add—and condition on—a new region C that is a real confounder. In the opposite case, an edge between A and B will be inferred if we do not include the real confounder C in the variable set. Given this dependency, connectivity results should be reported and interpreted relative to the variables considered. Inferring causal connectivity networks in the presence of unmeasured confounders, also known in the literature as latent confounders, is an ongoing research problem for which sophisticated methods have shown ways to model the presence of unmeasured confounders under strict assumptions. For example, the Fast Causal Inference (FCI) algorithm and its variants (Colombo et al., 2011; Malinsky & Spirtes, 2018; Spirtes et al., 1995) and algorithms based on independent component analysis (ICA) (Hoyer et al., 2006; Sanchez-Romero et al., 2018).

We have demonstrated that, despite these limitations, combinedFC is substantially more accurate than either bivariate correlation or partial correlation alone. We therefore recommend use of combinedFC in place of bivariate correlation or partial correlation in ongoing functional connectivity research. Notably, some other current methods might be just as valid (or even more so), but the complexity of those methods is problematic since researchers should not apply methods they do not understand well. It will therefore be critical for future research to develop causally valid methods that are easily comprehensible to researchers, in addition to providing clear explanations (especially regarding assumptions of methods) that aid in valid use of such methods by researchers.

## Acknowledgements

We would like to thank Clark Glymour for helpful conversations related to the ideas discussed here, and Takuya Ito for helpful discussions and data analysis support. The authors acknowledge support by the US National Institutes of Health under awards R01 AG055556 and R01 MH109520. Data were provided, in part, by the Human Connectome Project, WU-Minn Consortium (Principal Investigators: D. Van Essen and K. Ugurbil; 1U54MH091657) funded by the 16 NIH Institutes and Centers that support the NIH Blueprint for Neuroscience Research; and by the McDonnell Center for Systems Neuroscience at Washington University. The content is solely the responsibility of the authors and does not necessarily represent the official views of any of the funding agencies.

